# RAPID-DASH: Fast and Efficient Assembly of Guide RNA Arrays for Multiplexed CRISPR-Cas9 Applications

**DOI:** 10.1101/2025.04.09.648054

**Authors:** Asfar Lathif Salaudeen, Nicholas Mateyko, Carl G. de Boer

## Abstract

Guide RNA (gRNA) arrays can enable targeting multiple genomic loci simultaneously using CRISPR-Cas9. In this study, we present a streamlined and efficient method to rapidly construct gRNA arrays with up to 10 gRNA units in a single day. We demonstrate that gRNA arrays maintain robust functional activity across all positions, and can incorporate libraries of gRNAs, combining scalability and multiplexing. Our approach will streamline combinatorial perturbation research by enabling the economical and rapid construction, testing, and iteration of gRNA arrays. To facilitate the adaptation of this approach, we have made a web tool to design oligo sequences necessary to assemble gRNA arrays.

## Introduction

The CRISPR-Cas9 system has revolutionized the field of genome engineering by enabling targeted modification of specific sequences. This system is typically divided into two components: the nuclease protein (Cas9) and a guide RNA (gRNA) that directs Cas9 to its target region^1,2^. The spacer sequence of the gRNA can be customized to target regions of interest^4^. The nuclease domain has been customized in a variety of ways for different applications, including gene knockouts/knock-ins through double stranded breaks^3,5,6^, transcriptional regulation (CRISPRa^7^ and CRISPRi^8^), and rewriting the genome (base editors^9^ and prime editors^10^) and epigenome (CRISPRoff and CRISPRon)^11^. Concurrent delivery of multiple gRNAs is essential to target multiple regions within a cell simultaneously^12^. For instance, Perturb-seq screens target multiple genes within a single cell to identify genetic interactions and reveal cellular pathways^13^; lineage tracing and event recording often edit multiple recorder modules simultaneously to record the history of a cell^14–16^; and targeting the same gene with multiple gRNAs can increase the effect of CRISPRi and CRISPRa in pooled CRISPR-screens^17^.

Despite the numerous applications requiring gRNA multiplexing, concurrent cloning and delivery of multiple guides still faces several challenges. An ideal solution would ensure that a single construct includes all the gRNAs of interest to prevent different cells getting different amounts of each gRNA, as would happen with delivery of individually cloned gRNAs. For transfection, this may also be more efficient as, for the same mass of DNA, the concentration and thus the activity of each guide is higher when cloned on the same plasmid as opposed to transfecting individually pooled gRNA expressing plasmids. Integrating guides into the genome is often required for sustained expression and consistent dosage across cells; here too, having all guides on a single construct can facilitate their simultaneous integration because only a single selectable marker is needed to ensure successful delivery. Several approaches have been reported to build multiplexed gRNA systems^12^ (Table 1). Polycistronic pre-gRNAs can be processed into functioning gRNAs by simultaneous expression of the Csy4 nuclease^18–20^. However, this approach requires co-expression of exogenous Cys4 ribonuclease and gRNAs towards the end of the long RNA Polymerase III-transcribed polycistronic array were not functional^20^. It is also possible to leverage endogenous tRNA processing by interspersing gRNA units with tRNA sequences in a polycistronic pre-gRNA^21–24^, but this can perturb endogenous tRNA pools. Another technique involves creating arrays of self-contained gRNA units using Gibson assembly, but this approach tends to be inefficient, especially for many gRNA units, resulting in incomplete assemblies. This approach also requires multiple long primers for every gRNA units as the spacers sequences acts as the homology arms to mediate Gibson assembly^25^. Another study used Golden Gate assembly of cloned gRNA fragments into a single vector, including up to ten gRNAs total, but the initial gRNA cloning and sequence verification adds substantially to the total cloning time and cost^26^.

**Table 1:** Summary of RAPID-DASH compared to other gRNA array approaches.

| Study | Assembly approach | Maximum gRNAs assembled | gRNA processing method | Overall assembly time | Pre-cloning step | Scalability | Oligos per gRNA (length) | Other limitations |
| --- | --- | --- | --- | --- | --- | --- | --- | --- |
| RAPID-DASH | Golden gate assembly | 10 gRNAs (up to 20 gRNA units can be assembled using RAPID-DASH online tool) | Individual expression cassettes | 3 days | No | Scalable as gRNA oligos can be pooled during PCA to assemble gRNA array libraries | 1 (59 bp) | Long read sequencing required to screen clones |
| Vad-Nielsen et al.<sup>26</sup> | Golden gate assembly | 30 gRNAs (Assembled in 2 stages) | Individual expression cassettes | > 7 days | Yes | Cloning each gRNA restricts scalability | 2 (25 bp) | Labor intensive: multiple rounds of cloning, culturing and sequence validation |
| McCarty et al.<sup>18</sup> | Golden gate assembly | 12 gRNAs | Csy4 processing | > 2 days | No | Scalable as the gRNA units can be pooled before adding Type IIS overhangs for the array assembly (Not demonstrated in the study). | 1 (53 bp) | Labour intensive: multiple rounds of PCR, digestion and gel purifications. Requires Csy4 endonuclease for individual gRNA processing. |
| Kurata et al.<sup>20</sup> | Golden gate assembly | 10 gRNAs | Csy4 processing | > 5 days | Yes | Cloning each gRNA restricts scalability | 2 (25 bp) | Labor intensive: multiple rounds of cloning, culturing and sequence validation. Csy4 endonuclease for individual gRNA processing. |
| Zhang et al.<sup>23</sup> | Golden gate assembly | 8 gRNAs | tRNA processing | >2 days | No | gRNA spacer sequences being the Golden Gate overhangs restricts scalability. | 2 (44 to 60 bp) | The gRNA maturation could interfere with endogenous tRNA pool |
| Yuan et al.<sup>24</sup> | Golden gate assembly | 10 gRNAs | tRNA/Csy4/Ribozyme processing | 3-5 days | No | gRNA spacer sequences being the Golden Gate overhangs restricts scalability. | 2 (27 to 52 bp) | The gRNA maturation could interfere with endogenous tRNA pool (tRNA processing) or requires Csy4 endonuclease expression |
| Breunig et al.<sup>25</sup> | Gibson assembly | 8 gRNAs | Individual expression cassettes | 3 days | No | gRNA spacer sequences mediate homology for Gibson assembly thus restricting scalability. | 2 (36-45 bp) | Low assembly efficiency (34% for 4 gRNA assembly) |

<sup>#</sup>Scalability indicates using the approach to assemble multiple gRNA arrays or libraries of arrays simultaneously.

A previous study^27^ introduced a method to assemble multiple gRNAs into a single array using polymerase cycling assembly (PCA)^28^ to generate the gRNA units, which are then assembled in a specific order and cloned into the vector using Golden Gate assembly^29^ (**Fig. 1**). Using this approach, the authors assembled sets of 5 gRNAs into a single plasmid for multiplexed CRISPR-activation. Although this study demonstrated a modular way to assemble gRNA arrays, this method was not optimized or evaluated for assembly efficiency, error rates, or potential for scalable gRNA library generation. Building on this foundation, we optimized the approach into RAPID-DASH – **R**apid **A**ssembly of **P**CA-produced **I**ndividual **D**NAs and **D**irected **A**ssembly through typeII**S** over**H**angs for assembling arrays of gRNAs that enables at least 10 gRNAs to be cloned into an array within a single day. Each gRNA unit is created by PCA of a dsDNA U6 promoter, an ssDNA gRNA spacer oligo, and dsDNA gRNA scaffold terminator, resulting in a single dsDNA gRNA unit. A unique primer set is used for each position of the ordered gRNA array, which, in addition to amplifying the assembled gRNA units, adds Type IIS overhangs that enables ordered assembly of the gRNA units as an array (**Fig. 1a** step 1). During the Golden Gate guide assembly process, BsaI, a type IIS restriction enzyme, digests the ends of each gRNA unit, exposing overhangs that are complementary only to the adjacent gRNA unit, facilitating correct orderly assembly by ligation. (**Fig 1a** step 2). The lacZ gene within the destination plasmid enables efficient identification of colonies with the gRNA array via blue-white colony screening^30^ (**Supplementary Fig 1)**. In addition to cloning more gRNAs using RAPID-DASH, we provide a detailed characterization of the method, including measurement of assembly efficiency, quantitative error-rate profiling, and assessment of gRNA array library generation. These optimizations substantially enhance robustness, reduce synthesis costs, and establish RAPID-DASH as a versatile framework for constructing large, high-fidelity gRNA arrays suitable for multiplexed genome engineering applications.

**Figure 1:**
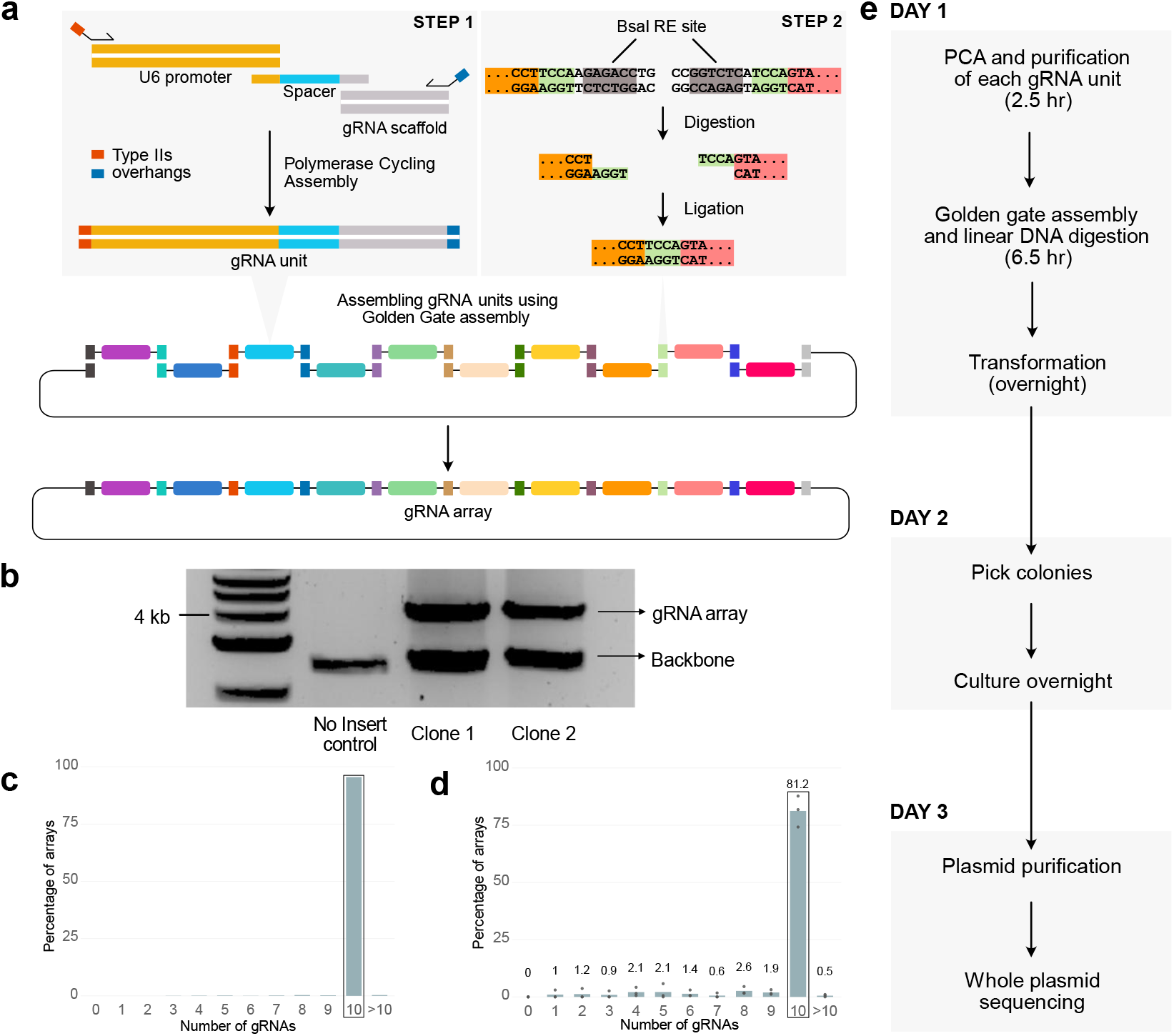
gRNA array assembly by RAPID-DASH. a, Overview of RAPID-DASH. gRNA units are assembled using PCA with unique primer sets that add Type IIS restriction enzyme recognition sites and 4 bp sequences that, when digested by BsaI, enable orderly assembly via Golden Gate assembly. The order of the assembly in this overview is determined by the color-coded Type IIS restriction site flanking the gRNA expression units. b, Screening bacterial clones by digesting out the assembled arrays. Gel electrophoresis image shows bands at ∼4kb, which is the expected size for 10 gRNA arrays. c,d, Bar plot showing the percentage of arrays with different number of gRNAs assembled from (c) a single clone whole plasmid and (d) bulk plasmid sequencing. Highlighted bar shows 10 gRNA arrays. Numbers above bars indicate average values. e, RAPID-DASH timeline.

## Methods

### gRNA unit generation using polymerase cycling assembly

Ten individual gRNA units were constructed by assembling double stranded U6 promoter, single stranded spacer sequence, and double stranded terminator scaffold sequence into a single unit. The U6 promoter and gRNA terminator scaffold fragments were amplified from a commonly used gRNA expression vector (MLM3636). The spacer sequence was ordered from IDT as a single stranded DNA oligo that included homology sequences to the U6 promoter and the terminator scaffold fragments. The assembly is mediated by a forward primer that binds to the U6 promoter and a reverse primer that binds to the terminator scaffold fragments. These primers were designed to include type II restriction enzyme (BsaI) overhangs that enable ordered assembly of the gRNA units into an array (**Supplementary Data**). We used the NEBridge Ligation fidelity GetSet tool to generate these type II overhangs (BsaI-HFv2 37-16 cycling). The sticky ends and the spacer oligos needed for assembling the gRNA units can be generated using our Shiny app: https://deboerlab.shinyapps.io/OligoDesigner/. The PCA reactions were set up as follows: 1 ul U6 promoter (5 ng), 1 ul gRNA terminator scaffold (5 ng), 2 ul spacer oligo (1 mM), 1.25 ul Forward Primers (10 uM), 1.25 ul Reverse Primers (10 uM), 25 ul Phusion High-Fidelity PCR Master Mix (Thermo F531), and nuclease free water to a total volume of 50 ul. The following setting were used in thermocycler: 98 C for 3 minutes, 35 cycles of 98 C for 10 seconds, 52 C for 30 seconds and 72 C for 12 seconds which was followed by a final 72 C for 8 minutes. The assembled units can be run on a 1% agarose gel to verify the length of units (433 bp). The gRNA units were then purified using AMPure XP beads. The PCA products were incubated with 1:1 ratio of AMPure XP beads and incubated for 5 minutes at room temperature. Following this, the product was placed on a magnetic separation rack and let the beads separate from the supernatant (∼ 3 minutes). The supernatant was then removed from the tubes and the beads were rinsed with 80% ethanol twice. The beads were then air dried for maximum 5 minutes to remove any residual ethanol from washing. The tubes were taken of the magnetic rack and the beads were resuspended in 20 ul of warm nuclease-free water and incubated at room temperature for 5 minutes. The tubes were placed back on to the magnetic rack and the PCA products are eluted from the supernatant. This clean up step is critical for the high efficiency of Golden Gate assembly.

### gRNA array generation using golden gate assembly

The destination vector (pAL10) incorporating the Golden Gate cloning site into which the gRNA array is inserted was generated from pFUS-B10 vector from the TALEN assembly kit^37^. Golden Gate assembly for cloning gRNA arrays was set up as follows: 1 ul destination vector (pAL10) (100 ng), 1 ul each of the purified gRNA units (50 ng), 1 ul of T4 DNA ligase (2000 U/ul, NEB M0202T), 1.5 ul FastDigest Eco31I (Thermo FD0293) (BsaI isoschizomer), 2 ul of 10X T4 DNA ligase buffer (NEB), and nuclease free water up to 20 ul. The following setting was used in the thermocycler: 5 minutes at 37 C and 5 minutes at 16 C for 30 cycles followed by 37 C for 10 minutes. The restriction enzyme was then inactivated at 75 C for 10 minutes. The reaction can be held at 4 C at this point. The reaction is mixed with 1 ul Plasmid-Safe ATP-Dependent DNase (Biosearch Technologies E3101K) and 1 ul of 25 mM ATP, and incubated at 37 C for 1 hour. This step was crucial to avoid recombination of linear DNA after transformation in bacterial cells^26^. 2 ul of the assembled product was used to transform NEB stable chemical competent cells using the manufacturer-supplied heat-shock transformation protocol and plated on LB agar with 50 ug/ml spectinomycin along with IPTG and X-gal for blue/white screening^30^. The transformants were incubated at 30 C for 16 hr. Plasmids isolated from white colonies were screened for correct gRNA array assembly via BsmB1-v2 digestion (NEB R0739, Manufacturer’s protocol). For bulk plasmid sequencing, the transformants were directly inoculated into LB media with 50 ug/ml spectinomycin. Whole plasmid sequencing to validate the arrays was performed by Plasmidsaurus using Oxford Nanopore Technology.

The probability of picking a colony that is suitable for downstream use is estimated as follows. Mutations within the PCA primer regions are ignored in the following calculations as they are most likely to be inconsequential. All mutation indicated below only refer to mutations observed within either the U6 promoter, gRNA spacer or the gRNA scaffold of a gRNA unit.

The probability of a gRNA unit having no mutations is given by:

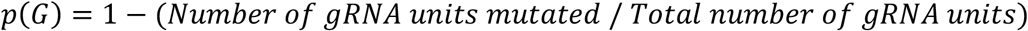

Based on our data, *p*(*G*) = 92.5%.

The probability of an array with *k* gRNA units having with no mutations, which assumes independence between gRNA units (they are synthesized separately) is given by:

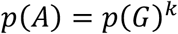

Our data suggest that the probability of an assembly being full length, *p*(*F*), is 81%. This is assuming each of the nanopore reads from bulk plasmid sequencing stems from an individual clone and the assembly efficiency is the proportion of nanopore reads that are the full-length assembly.

The probability of a picked clone being a full-length assembly and having no mutations in any gRNA unit is given by:

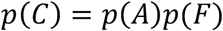

The probability of picking at least one usable clone (full-length assembly with no mutations) out of *n* colonies (assuming each is an independent transformation event) is given by:

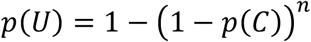

### GFP reporter cell line transfection

To validate the array, we used a mutated GFP reporter HEK293T cell line from Sakata et al, 2020^31^. gRNA arrays consisting of the GFP targeting gRNA within each position of the array were generated as above. We also included a gRNA array that did not include the GFP targeting gRNA and plasmids that expressed individual gRNAs as controls. For the GFP reporter assay, 1 × 10^5^ cells were plated in wells of a 48-well plate the day before transfection. Cells were transfected with equimolar (40 fmoles) Target-AID base editor and either the single GFP targeting gRNA plasmid or gRNA arrays in respective wells. Transporter5 reagent (Polysciences 26008) was used for transfecting the DNA into the cells using manufacturer’s protocol. Transfection Media was replaced with fresh media 24 hrs after transfection. Cells were harvested 72 hrs after transfection for flow cytometry analysis. Three independent replicates were performed for each transfection.

### Flow cytometry analysis

Cells were analyzed for GFP expression using Cytoflex LX Analyser and gated using FlowJo (v10) (**Supplementary Fig 2**). Singlet cells were extracted using FlowJo (v10) and then further processed using tidyverse (v2.0.0) in R 4.4.2.

### gRNA array library generation

To generate gRNA arrays with gRNAs randomly sampled from a pool, we pooled the ten single stranded spacer oligos in equimolar ratio prior to gRNA unit generation via PCA to mimic commercially synthesized oligo pools. gRNA units and the subsequent assembly were done as described above. Bulk plasmid sequencing was performed by Plasmidsaurus using Oxford Nanopore Technology.

### Nanopore Sequencing Analysis

The reads in fastq files from nanopore sequencing often contain the whole plasmid sequences. gRNA array sequences from the whole plasmid sequences were extracted using the ‘get_plasmid_inserts’ custom function which extracts the gRNA array inserts based on input flanking sequences in the backbone. To gauge gRNA assembly efficiency, the extracted inserts were binned by length into the number of gRNA units each represents based on the known length of each gRNA unit (**Supplementary Table 2)**. To get the distribution of gRNA sequences within the randomized gRNA array library, we filtered the nanopore reads to full-length gRNA arrays with 10 gRNA units and used the RapidFuzz alignment^38^ approach to map each gRNA sequence to the known sequences within the gRNA pool. The resultant gRNA vs position map was then plotted using ggplot2 (v3.5.1) in R 4.4.2. Detailed analysis can be found in our GitHub repository.

### Statistical Analysis

Two-sample Welch’s t-test was performed on the GFP reporter flow cytometry data to compare the GFP reporter activation efficiency of each of the GFP targeting guide in 10 guide positions to the non-targeting control and to the single GFP targeting guide. Bonferroni’s method was used to adjust for multiple hypothesis testing within each comparison set (vs. NTC and vs. GFP-only). All the statistical analysis were performed in R 4.4.2. Detailed analysis can be found in our GitHub repository.

## Results and Discussion

We tested RAPID-DASH to assemble an array of 10 gRNAs (**Methods**). Digestion of plasmids isolated from individual clones revealed that the assembled arrays are of expected size (4kb) (**Fig. 1b**). To verify the sequence of the assembled gRNA arrays, we extracted plasmid from a single clone and sequenced it using nanopore sequencing. We verified that it included all 10 gRNAs (**Fig. 1c**), and sequences were correct (**Supplementary File 2**) and assembled in the expected order. Whole plasmid sequencing of a pool of transformants (i.e. plasmids extracted from the recovered transformation mix grown in liquid LB + Spectinomycin, without plating) revealed that, on average, 81% (n=3 replicates) of the assemblies have all 10 gRNA units (**Fig. 1d, Supplementary Table 1**). In order to estimate the error-rate within the assembly, we sequenced 28 gRNA array clones using nanopore sequencing, breaking it down into mutations that are presumably inconsequential as they are only used for gRNA unit amplification during PCA, and potentially deleterious mutations impacting the gRNA (U6 promoter, gRNA spacer and scaffold), finding that 7.5% of gRNA units had one or more potentially deleterious mutation (**Supplementary Figure 4**). Given these results, screening 4 colonies has an 84% chance of recovering a full-length array with no putatively deleterious mutations within any gRNA unit. (**Methods**). Pre-screening the colonies by restriction digest can help eliminate incomplete assemblies (**Fig. 1b**). Further, while we did not purify our oligos (standard desalting only), this would likely substantially reduce the error rate, although at a high cost per oligo. Depending on a lab’s individual circumstances, these may or may not be more economical than just directly screening colonies.

To validate the functionality of gRNA units within the array, we used a cell line that can report successful CRISPR base editing by activating a GFP gene^31^. Here, the GFP gene is initially defective because its start codon has been mutated to GTG. By targeting the Target-AID base editor^32^ to the start codon with a gRNA, the start codon is repaired to an ATG, enabling gRNA activity to be assayed easily via cells turning green (**Fig 2a**). We cloned the GFP targeting gRNA within each position of the gRNA arrays independently and observed that the 10 gRNA positions had similar efficacy, activating GFP in 20.17% to 26.53% of cells (**Fig 2b)**, which was far more than the non-targeting gRNA control (0.48% GFP positive; Welch’s t-test p < 0.01 for all gRNAs; **Supplementary Table 3**). The efficacy of gRNAs in arrays was slightly less than a plasmid containing only a single GFP-targeting gRNA (30.56% GFP positive; 0.0218 ≤ p ≤ 0.419 for all gRNAs; **Supplementary Table 3**), which was expected due to the higher transfection efficiency of the smaller GFP-only construct. These results illustrate that these gRNA arrays are suitable for multiplexed gene editing as all the positions within the array are functional and have similar editing efficiencies.

**Figure 2:**
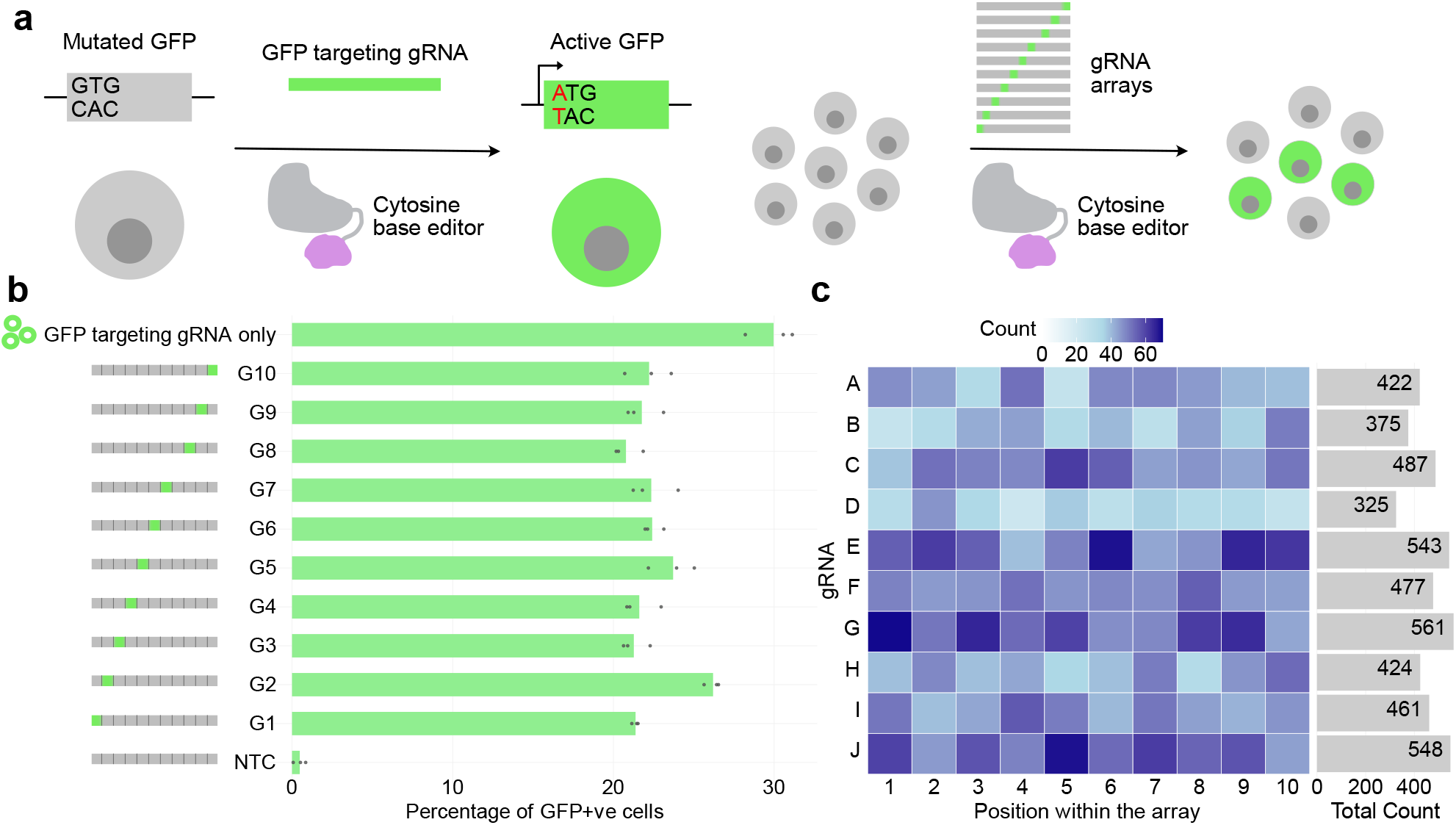
Functional validation of the assembled arrays. a, GFP reporter assay for gRNA array validation. b, Bar chart showing the percentage of the GFP reporter cells (x axis) activated by gRNA arrays with GFP-targeting gRNA at each position (y axis) within the array. NTC=non-targeting control with no GFP-targeting guide c, Abundance (colour) of each guide RNA (gRNA) sequences (y axis) across different positions in an array (x axis) within the nanopore reads of a bulk plasmid sequence when using a pooled gRNA sequences during PCA for each array position. The bar plot on the right depicts the total count for each gRNA across all the ten positions within the array.

For applications requiring a randomized combinatorial genome editing or perturbation, it may be desirable to create libraries of gRNA arrays, each with gRNAs targeting distinct genomic loci. For example, in a large-scale CRISPRa/CRISPRi functional genomics screen, one might aim to interrogate the regulatory effect of ∼100,000 genomic elements (e.g., promoters or enhancers) by randomly activating or repressing 10 elements per cell. This enables combinatorial perturbation, where different cells receive different combinations of targets, allowing higher-order interaction effects to be observed. By using pools of gRNA sequences during the PCA step, such libraries can be created with our approach. As a proof of concept, we tested this by including 10 gRNA sequences for each of the gRNA unit PCAs, and then assembled them as before into a library, where each library member gets an array of 10 gRNAs, each randomly sampled from the 10 possibilities. For most applications, one would want to sample from a larger pool of gRNAs, but for this demonstration we limited it to 10 so that we could obtain better sequencing coverage, although this strategy nonetheless could produce 10 billion (10^10^) unique correctly assembled arrays. Nanopore sequencing of these pooled arrays showed all 10 gRNAs were present in all 10 positions (**Fig 2c**). Combined with the relatively low cost per oligo when ordering pooled oligos, this approach can be scaled up to include many more gRNA sequences to create libraries of gRNA arrays efficiently, enabling low-cost multiplexed perturbation assays. Of note, because the PCA of each gRNA unit is performed separately, one could limit the possible gRNAs in each position, ensuring that only certain combinations of gRNAs are created.

Using RAPID-DASH, we can generate and validate gRNA arrays with only a single day of hands-on time (**Fig. 1e**). RAPID-DASH offers substantial cost savings because only a short (∼60nt) spacer sequence needs to be ordered for each new gRNA. Researchers can also skip the PCA step if they have gRNA units synthesized (433 bp; e.g., gBlocks), however, this will increase the cost substantially as the PCA-based synthesis requires only a single 59 nt spacer oligo for each new gRNA unit. Further, the high-efficiency of our approach saves additional time and labor, reducing the need for layers of screening and enabling one to use libraries of gRNA arrays directly in downstream applications. The major bottlenecks of RAPID-DASH reflect oligo synthesis and sequencing.

Although we only scale to 10 gRNA units, which should satisfy most applications for the time being, we anticipate that RAPID-DASH can be scaled further by using appropriate Type II restriction enzyme overhangs to assemble up to 52 gRNA units, although this would result in a plasmid of substantial size (∼24kb) and likely reduce the efficiency of the assembly and stability of the assembled arrays RAPID-DASH enables faster and more efficient construction and more robust delivery of gRNA arrays, facilitating rapid experimental iteration at scale (Table 1). In order to facilitate adoption of RAPID-DASH, we developed a Shiny app (https://deboerlab.shinyapps.io/OligoDesigner/) that enables design of spacer sequence oligos for assembly of up to 20 gRNA units within a single array. Arrays assembled using RAPID-DASH can be transfected directly, used to create virus-like particles containing CRISPR ribonucleoproteins^33^, or cloned into PiggyBac vectors furthering the scale to perform multiplexed CRISPR screens^34^. Readouts of CRISPR screens vary by experiment, but we anticipate that RAPID-DASH will be most useful for approaches that read out the mutations directly^35^, incorporate a barcode that can be used to infer the guide array^36^, or via directly capturing and sequencing gRNAs (10X CRISPR Guide Capture)^17^, where imputation could be used to infer gRNA presence even when unobserved if it was known to be on the same array.

## Material Availability

Destination vector plasmid (pAL10) used to clone the gRNA arrays is deposited to Addgene for non-commercial use (Addgene #236006).

## Data Availability

Raw nanopore sequencing of assembled arrays are available at https://github.com/de-Boer-Lab/RAPID-DASH/. Code used for processing nanopore data and plotting are available at the https://github.com/de-Boer-Lab/RAPID-DASH/

## Supporting Information

The supplementary materials (.docx) file contains Blue-white screening of bacterial colonies following gRNA array assembly (Supplementary Figure 1), gating summary for flow cytometry data (Supplementary Figure 2), example mutations seen in gRNA arrays (Supplementary Figure 3), percentage of gRNA units mutated within the screened arrays (Supplementary Figure 4), gRNA array assembly efficiency by in percentage from bulk plasmid sequencing (Supplementary Table 1), read length binning thresholds for calculating the number of gRNAs in an array (Supplementary Table 2), pairwise comparisons of GFP reporter activation for each gRNA position within the array versus NTC and GFP-only controls (Supplementary Table 3) and the oligos and sequences for gRNA assembly. Supplementary File 2 (.pdf) contains the alignment of a full-length clonal gRNA array plasmid sequence generated via Nanopore sequencing to the desired sequence.

## Abbreviations

CRISPR: Clustered Regularly Interspaced Short Palindromic Repeats
gRNA: Guide RNA
RAPID-DASH: Rapid Assembly of PCA-produced Individual DNAs and Directed Assembly through typeIIS overHangs
PCA: Polymerase Cycling Assembly
dsDNA: Double stranded DNA
ssDNA: Single stranded DNA
GFP: Green Fluorescent Protein
NTC: Non-Targeting Control

## Author Information

### Author Contributions

RAPID-DASH idea was conceived by A.L.S and C.G.D. A.L.S performed the experiments and analyzed the data. A.L.S and N.M wrote the code for nanopore sequencing data analysis. A.L.S, and C.G.D. wrote the manuscript.

## Acknowledgments

We thank Dr. Nozomu Yachie (University of British Columbia) for providing us with the GFP reporter cell line used in this study. Research reported in this publication was supported by the Canadian Institutes for Health Research (PJT-180537), the National Science and Engineering Research Council of Canada (RGPIN-2020-05425), and the National Cancer Institute of the National Institutes of Health under award number R01CA279795. A.L.S and N.M. were supported by a UBC 4-Year Fellowship. C.G.D. is a Michael Smith Health Research BC Scholar. The content is solely the responsibility of the authors and does not necessarily represent the official views of the funders

## References

1. Mojica, F. J. M., Díez-Villaseñor, C., García-Martínez, J. & Soria, E. Intervening sequences of regularly spaced prokaryotic repeats derive from foreign genetic elements. J. Mol. Evol. 60, 174–182 (2005).

2. Pourcel, C., Salvignol, G. & Vergnaud, G. CRISPR elements in Yersinia pestis acquire new repeats by preferential uptake of bacteriophage DNA, and provide additional tools for evolutionary studies. Microbiol. Read. Engl. 151, 653–663 (2005).

3. Jinek, M. et al. A programmable dual-RNA-guided DNA endonuclease in adaptive bacterial immunity. Science 337, 816–821 (2012).

4. Doudna, J. A. & Charpentier, E. The new frontier of genome engineering with CRISPR-Cas9. Science 346, 1258096 (2014).

5. Mali, P. et al. RNA-Guided Human Genome Engineering via Cas9. Science 339, 823–826 (2013).

6. Cong, L. et al. Multiplex Genome Engineering Using CRISPR/Cas Systems. Science 339, 819–823 (2013).

7. Gilbert, L. A. et al. CRISPR-Mediated Modular RNA-Guided Regulation of Transcription in Eukaryotes. Cell 154, 442–451 (2013).

8. Maeder, M. L. et al. CRISPR RNA–guided activation of endogenous human genes. Nat. Methods 10, 977–979 (2013).

9. Komor, A. C., Kim, Y. B., Packer, M. S., Zuris, J. A. & Liu, D. R. Programmable editing of a target base in genomic DNA without double-stranded DNA cleavage. Nature 533, 420–424 (2016).

10. Anzalone, A. V. et al. Search-and-replace genome editing without double-strand breaks or donor DNA. Nature 576, 149–157 (2019).

11. Nuñez, J. K. et al. Genome-wide programmable transcriptional memory by CRISPR-based epigenome editing. Cell 184, 2503-2519.e17 (2021).

12. McCarty, N. S., Graham, A. E., Studená, L. & Ledesma-Amaro, R. Multiplexed CRISPR technologies for gene editing and transcriptional regulation. Nat. Commun. 11, 1281 (2020).

13. Adamson, B. et al. A Multiplexed Single-Cell CRISPR Screening Platform Enables Systematic Dissection of the Unfolded Protein Response. Cell 167, 1867-1882.e21 (2016).

14. Sheth, R. U. & Wang, H. H. DNA-based memory devices for recording cellular events. Nat. Rev. Genet. 19, 718–732 (2018).

15. Tang, W. & Liu, D. R. Rewritable multi-event analog recording in bacterial and mammalian cells. Science 360, eaap8992 (2018).

16. Sheth, R. U., Yim, S. S., Wu, F. L. & Wang, H. H. Multiplex recording of cellular events over time on CRISPR biological tape. Science 358, 1457–1461 (2017).

17. Replogle, J. M. et al. Combinatorial single-cell CRISPR screens by direct guide RNA capture and targeted sequencing. Nat. Biotechnol. 38, 954–961 (2020).

18. McCarty, N. S., Shaw, W. M., Ellis, T. & Ledesma-Amaro, R. Rapid Assembly of gRNA Arrays via Modular Cloning in Yeast. ACS Synth. Biol. 8, 906–910 (2019).

19. Ferreira, R., Skrekas, C., Nielsen, J. & David, F. Multiplexed CRISPR/Cas9 Genome Editing and Gene Regulation Using Csy4 in Saccharomyces cerevisiae. ACS Synth. Biol. 7, 10–15 (2018).

20. Kurata, M. et al. Highly multiplexed genome engineering using CRISPR/Cas9 gRNA arrays. PLoS ONE 13, e0198714 (2018).

21. Port, F. & Bullock, S. L. Augmenting CRISPR applications in Drosophila with tRNA-flanked Cas9 and Cpf1 sgRNAs. Nat. Methods 13, 852–854 (2016).

22. Xie, K., Minkenberg, B. & Yang, Y. Boosting CRISPR/Cas9 multiplex editing capability with the endogenous tRNA-processing system. Proc. Natl. Acad. Sci. U. S. A. 112, 3570–3575 (2015).

23. Zhang, Y. et al. A gRNA-tRNA array for CRISPR-Cas9 based rapid multiplexed genome editing in Saccharomyces cerevisiae. Nat. Commun. 10, 1053 (2019).

24. Yuan, G., Martin, S., Hassan, M. M., Tuskan, G. A. & Yang, X. PARA: A New Platform for the Rapid Assembly of gRNA Arrays for Multiplexed CRISPR Technologies. Cells 11, 2467 (2022).

25. Breunig, C. T. et al. One step generation of customizable gRNA vectors for multiplex CRISPR approaches through string assembly gRNA cloning (STAgR). PLOS ONE 13, e0196015 (2018).

26. Vad-Nielsen, J., Lin, L., Bolund, L., Nielsen, A. L. & Luo, Y. Golden Gate Assembly of CRISPR gRNA expression array for simultaneously targeting multiple genes. Cell. Mol. Life Sci. 73, 4315–4325 (2016).

27. Balboa, D. et al. Conditionally Stabilized dCas9 Activator for Controlling Gene Expression in Human Cell Reprogramming and Differentiation. Stem Cell Rep. 5, 448–459 (2015).

28. TerMaat, J. R., Pienaar, E., Whitney, S. E., Mamedov, T. G. & Subramanian, A. Gene synthesis by integrated polymerase chain assembly and PCR amplification using a high-speed thermocycler. J. Microbiol. Methods 79, 295–300 (2009).

29. Engler, C., Kandzia, R. & Marillonnet, S. A One Pot, One Step, Precision Cloning Method with High Throughput Capability. PLoS ONE 3, e3647 (2008).

30. Green, M. R. & Sambrook, J. Screening Bacterial Colonies Using X-Gal and IPTG: α-Complementation. Cold Spring Harb. Protoc. 2019, (2019).

31. Sakata, R. C. et al. Base editors for simultaneous introduction of C-to-T and A-to-G mutations. Nat. Biotechnol. 38, 865–869 (2020).

32. Nishida, K. et al. Targeted nucleotide editing using hybrid prokaryotic and vertebrate adaptive immune systems. Science 353, aaf8729 (2016).

33. Banskota, S. et al. Engineered virus-like particles for efficient in vivo delivery of therapeutic proteins. Cell 185, 250-265.e16 (2022).

34. Chardon, F. M. et al. Multiplex, single-cell CRISPRa screening for cell type specific regulatory elements. Nat. Commun. 15, 8209 (2024).

35. Martyn, G. E. et al. Rewriting regulatory DNA to dissect and reprogram gene expression. 2023.12.20.572268 Preprint at 10.1101/2023.12.20.572268 (2023).

36. Wong, A. S. L. et al. Multiplexed barcoded CRISPR-Cas9 screening enabled by CombiGEM. Proc. Natl. Acad. Sci. 113, 2544–2549 (2016).

37. Cermak, T. et al. Efficient design and assembly of custom TALEN and other TAL effector-based constructs for DNA targeting. Nucleic Acids Res. 39, e82 (2011).

38. Max Bachmann. rapidfuzz/RapidFuzz: Release 3.8.1. Zenodo 10.5281/ZENODO.10938887 (2024).

