## SupplementaryFile2 for "RAPID-DASH: Fast and Efficient Assembly of Guide RNA Arrays for Multiplexed CRISPR-Cas9 Applications"

Page 1 Consensus Sequence from Nanopore Sequencing

```
               1                                                                                82
Template       ttgagatcctttttttctgcgcgtaatctgctgcttgcaaacaaaaaaaccaccgctaccagcggtggtttgtttgccggat
Clonal Se...   TTGAGATCCTTTTTTTCTGCGCGTAATCTGCTGCTTGCAAACAAAAAAACCACCGCTACCAGCGGTGGTTTGTTTGCCGGAT

..................................................................................................

               83                                                                              164
Template       caagagctaccaactctttttccgaaggtaactggcttcagcagagcgcagataccaaatactgttcttctagtgtagccgt
Clonal Se...   CAAGAGCTACCAACTCTTTTTCCGAAGGTAACTGGCTTCAGCAGAGCGCAGATACCAAATACTGTTCTTCTAGTGTAGCCGT

..................................................................................................

               165                                                                             246
Template       agttaggccaccacttcaagaactctgtagcaccgcctacatacctcgctctgctaatcctgttaccagtggctgctgccag
Clonal Se...   AGTTAGGCCACCACTTCAAGAACTCTGTAGCACCGCCTACATACCTCGCTCTGCTAATCCTGTTACCAGTGGCTGCTGCCAG

..................................................................................................

               247                                                                             328
Template       tggcgataagtcgtgtcttaccgggttggactcaagacgatagttaccggataaggcgcagcggtcgggctgaacggggggt
Clonal Se...   TGGCGATAAGTCGTGTCTTACCGGGTTGGACTCAAGACGATAGTTACCGGATAAGGCGCAGCGGTCGGGCTGAACGGGGGGT

..................................................................................................

               329                                                                             410
Template       tcgtgcacacagcccagcttggagcgaacgacctacaccgaactgagatacctacagcgtgagctatgagaaagcgccacgc
Clonal Se...   TCGTGCACACAGCCCAGCTTGGAGCGAACGACCTACACCGAACTGAGATACCTACAGCGTGAGCTATGAGAAAGCGCCACGC

..................................................................................................

               411                                                                             492
Template       ttcccgaagggagaaaggcggacaggtatccggtaagcggcagggtcggaacaggagagcgcacgagggagcttccaggggg
Clonal Se...   TTCCCGAAGGGAGAAAGGCGGACAGGTATCCGGTAAGCGGCAGGGTCGGAACAGGAGAGCGCACGAGGGAGCTTCCAGGGGG

..................................................................................................

               493                                                                             574
Template       aaacgcctggtatctttatagtcctgtcgggtttcgccacctctgacttgagcgtcgatttttgtgatgctcgtcagggggg
Clonal Se...   AAACGCCTGGTATCTTTATAGTCCTGTCGGGTTTCGCCACCTCTGACTTGAGCGTCGATTTTTGTGATGCTCGTCAGGGGGG

..................................................................................................

               575                                                                             656
Template       cggagcctatggaaaaacgccagcaacgcggcctttttacggttcctggccttttgctggccttttgctcacatgttctttc
Clonal Se...   CGGAGCCTATGGAAAAACGCCAGCAACGCGGCCTTTTTACGGTTCCTGGCCTTTTGCTGGCCTTTTGCTCACATGTTCTTTC

..................................................................................................

               657                                                                             738
Template       ctgcgttatcccctgattctgtggataaccgtattaccgcctttgagtgagctgataccgctcgccgcagccgaacgaccga
Clonal Se...   CTGCGTTATCCCCTGATTCTGTGGATAACCGTATTACCGCCTTTGAGTGAGCTGATACCGCTCGCCGCAGCCGAACGACCGA

..................................................................................................

               739                                                                             820
Template       gcgcagcgagtcagtgagcgaggaagcggaagagcgcccaatacgcaaaccgcctctccccgcgcgttggccgattcattaa
Clonal Se...   GCGCAGCGAGTCAGTGAGCGAGGAAGCGGAAGAGCGCCCAATACGCAAACCGCCTCTCCCCGCGCGTTGGCCGATTCATTAA

..................................................................................................

               821                                                                             902
Template       tgcagctggcacgacaggtttcccgactggaaagcgggcagtgagcgcaacgcaattaatacgcgtaccgctagccaggaag
Clonal Se...   TGCAGCTGGCACGACAGGTTTCCCGACTGGAAAGCGGGCAGTGAGCGCAACGCAATTAATACGCGTACCGCTAGCCAGGAAG

..................................................................................................
```

```
              903                                                                             984
Template      agtttgtagaaacgcaaaaaggccatccgtcaggatggccttctgcttagtttgatgcctggcagtttatggcgggcgtcct
Clonal Se...  AGTTTGTAGAAACGCAAAAAGGCCATCCGTCAGGATGGCCTTCTGCTTAGTTTGATGCCTGGCAGTTTATGGCGGGCGTCCT

..................................................................................................

              985                                                                            1066
Template      gcccgccaccctccgggccgttgcttcacaacgttcaaatccgctcccggcggatttgtcctactcaggagagcgttcaccg
Clonal Se...  GCCCGCCACCCTCCGGGCCGTTGCTTCACAACGTTCAAATCCGCTCCCGGCGGATTTGTCCTACTCAGGAGAGCGTTCACCG

..................................................................................................

              1067                                                                           1148
Template      acaaacaacagataaaacgaaaggcccagtcttccgactgagcctttcgttttatttgatgcctggcagttccctactctcg
Clonal Se...  ACAAACAACAGATAAAACGAAAGGCCCAGTCTTCCGACTGAGCCTTTCGTTTTATTTGATGCCTGGCAGTTCCCTACTCTCG

..................................................................................................

              1149                                                                           1230
Template      cgttaacgctagcatggatgttttcccagtcacgacgttgtaaaacgacggccagtcttaagcgtctcatggcctgaccccg
Clonal Se...  CGTTAACGCTAGCATGGATGTTTTCCCAGTCACGACGTTGTAAAACGACGGCCAGTCTTAAGCGTCTCATGGCCTGACCCCG

..................................................................................................

              1231                                                                           1312
Template      gaccaagtggtggggtagtaaaacgacggccagtgagggcctatttcccatgattccttcatatttgcatatacgatacaag
Clonal Se...  GACCAAGTGGTGGGGTAGTAAAACGACGGCCAGTGAGGGCCTATTTCCCATGATTCCTTCATATTTGCATATACGATACAAG

..................................................................................................

              1313                                                                           1394
Template      gctgttagagagataattagaattaatttgactgtaaacacaaagatattagtacaaaatacgtgacgtagaaagtaataat
Clonal Se...  GCTGTTAGAGAGATAATTAGAATTAATTTGACTGTAAACACAAAGATATTAGTACAAAATACGTGACGTAGAAAGTAATAAT

..................................................................................................

              1395                                                                           1476
Template      ttcttgggtagtttgcagttttaaaattatgttttaaaatggactatcatatgcttaccgtaacttgaaagtatttcgattt
Clonal Se...  TTCTTGGGTAGTTTGCAGTTTTAAAATTATGTTTTAAAATGGACTATCATATGCTTACCGTAACTTGAAAGTATTTCGATTT

..................................................................................................

              1477                                                                           1558
Template      cttggctttatatatcttgtggaaaggacgaaacaccgggaatcccttctgcagcaccgttttagagctagaaatagcaagt
Clonal Se...  CTTGGCTTTATATATCTTGTGGAAAGGACGAAACACCGGGAATCCCTTCTGCAGCACCGTTTTAGAGCTAGAAATAGCAAGT

..................................................................................................

              1559                                                                           1640
Template      taaaataaggctagtccgttatcaacttgaaaaagtggcaccgagtcggtgctttttttcatggtcatagctgtttccttag
Clonal Se...  TAAAATAAGGCTAGTCCGTTATCAACTTGAAAAAGTGGCACCGAGTCGGTGCTTTTTTTCATGGTCATAGCTGTTTCCTTAG

..................................................................................................

              1641                                                                           1722
Template      agtaaaacgacggccagtgagggcctatttcccatgattccttcatatttgcatatacgatacaaggctgttagagagataa
Clonal Se...  AGTAAAACGACGGCCAGTGAGGGCCTATTTCCCATGATTCCTTCATATTTGCATATACGATACAAGGCTGTTAGAGAGATAA

..................................................................................................

              1723                                                                           1804
Template      ttagaattaatttgactgtaaacacaaagatattagtacaaaatacgtgacgtagaaagtaataatttcttgggtagtttgc
Clonal Se...  TTAGAATTAATTTGACTGTAAACACAAAGATATTAGTACAAAATACGTGACGTAGAAAGTAATAATTTCTTGGGTAGTTTGC

..................................................................................................
```

```
             1805                                                                           1886
Template     agttttaaaattatgttttaaaatggactatcatatgcttaccgtaacttgaaagtatttcgatttcttggctttatatatc
Clonal Se... AGTTTTAAAATTATGTTTTAAAATGGACTATCATATGCTTACCGTAACTTGAAAGTATTTCGATTTCTTGGCTTTATATATC

...............................................................................................

             1887                                                                           1968
Template     ttgtggaaaggacgaaacaccggagtccgagcagaagaagaagttttagagctagaaatagcaagttaaaataaggctagtc
Clonal Se... TTGTGGAAAGGACGAAACACCGGAGTCCGAGCAGAAGAAGAAGTTTTAGAGCTAGAAATAGCAAGTTAAAATAAGGCTAGTC

...............................................................................................

             1969                                                                           2050
Template     cgttatcaacttgaaaaagtggcaccgagtcggtgctttttttcatggtcatagctgtttcctctccgtaaaacgacggcca
Clonal Se... CGTTATCAACTTGAAAAAGTGGCACCGAGTCGGTGCTTTTTTTCATGGTCATAGCTGTTTCCTCTCCGTAAAACGACGGCCA

...............................................................................................

             2051                                                                           2132
Template     gtgagggcctatttcccatgattccttcatatttgcatatacgatacaaggctgttagagagataattagaattaatttgac
Clonal Se... GTGAGGGCCTATTTCCCATGATTCCTTCATATTTGCATATACGATACAAGGCTGTTAGAGAGATAATTAGAATTAATTTGAC

...............................................................................................

             2133                                                                           2214
Template     tgtaaacacaaagatattagtacaaaatacgtgacgtagaaagtaataatttcttgggtagtttgcagttttaaaattatgt
Clonal Se... TGTAAACACAAAGATATTAGTACAAAATACGTGACGTAGAAAGTAATAATTTCTTGGGTAGTTTGCAGTTTTAAAATTATGT

...............................................................................................

             2215                                                                           2296
Template     tttaaaatggactatcatatgcttaccgtaacttgaaagtatttcgatttcttggctttatatatcttgtggaaaggacgaa
Clonal Se... TTTAAAATGGACTATCATATGCTTACCGTAACTTGAAAGTATTTCGATTTCTTGGCTTTATATATCTTGTGGAAAGGACGAA

...............................................................................................

             2297                                                                           2378
Template     acaccgtttatcacaggctccaggaagttttagagctagaaatagcaagttaaaataaggctagtccgttatcaacttgaaa
Clonal Se... ACACCGTTTATCACAGGCTCCAGGAAGTTTTAGAGCTAGAAATAGCAAGTTAAAATAAGGCTAGTCCGTTATCAACTTGAAA

...............................................................................................

             2379                                                                           2460
Template     aagtggcaccgagtcggtgctttttttcatggtcatagctgtttcctatcagtaaaacgacggccagtgagggcctatttcc
Clonal Se... AAGTGGCACCGAGTCGGTGCTTTTTTTCATGGTCATAGCTGTTTCCTATCAGTAAAACGACGGCCAGTGAGGGCCTATTTCC

...............................................................................................

             2461                                                                           2542
Template     catgattccttcatatttgcatatacgatacaaggctgttagagagataattagaattaatttgactgtaaacacaaagata
Clonal Se... CATGATTCCTTCATATTTGCATATACGATACAAGGCTGTTAGAGAGATAATTAGAATTAATTTGACTGTAAACACAAAGATA

...............................................................................................

             2543                                                                           2624
Template     ttagtacaaaatacgtgacgtagaaagtaataatttcttgggtagtttgcagttttaaaattatgttttaaaatggactatc
Clonal Se... TTAGTACAAAATACGTGACGTAGAAAGTAATAATTTCTTGGGTAGTTTGCAGTTTTAAAATTATGTTTTAAAATGGACTATC

...............................................................................................

             2625                                                                           2706
Template     atatgcttaccgtaacttgaaagtatttcgatttcttggctttatatatcttgtggaaaggacgaaacaccgggcccagact
Clonal Se... ATATGCTTACCGTAACTTGAAAGTATTTCGATTTCTTGGCTTTATATATCTTGTGGAAAGGACGAAACACCGGGCCCAGACT

...............................................................................................
```

```
             2707                                                                            2788
Template     gagcacgtgagttttagagctagaaatagcaagttaaaataaggctagtccgttatcaacttgaaaaagtggcaccgagtcg
Clonal Se... GAGCACGTGAGTTTTAGAGCTAGAAATAGCAAGTTAAAATAAGGCTAGTCCGTTATCAACTTGAAAAAGTGGCACCGAGTCG

..................................................................................................

             2789                                                                            2870
Template     gtgctttttttcatggtcatagctgtttcctctgagtaaaacgacggccagtgagggcctatttcccatgattccttcatat
Clonal Se... GTGCTTTTTTTCATGGTCATAGCTGTTTCCTCTGAGTAAAACGACGGCCAGTGAGGGCCTATTTCCCATGATTCCTTCATAT

..................................................................................................

             2871                                                                            2952
Template     ttgcatatacgatacaaggctgttagagagataattagaattaatttgactgtaaacacaaagatattagtacaaaatacgt
Clonal Se... TTGCATATACGATACAAGGCTGTTAGAGAGATAATTAGAATTAATTTGACTGTAAACACAAAGATATTAGTACAAAATACGT

..................................................................................................

             2953                                                                            3034
Template     gacgtagaaagtaataatttcttgggtagtttgcagttttaaaattatgttttaaaatggactatcatatgcttaccgtaac
Clonal Se... GACGTAGAAAGTAATAATTTCTTGGGTAGTTTGCAGTTTTAAAATTATGTTTTAAAATGGACTATCATATGCTTACCGTAAC

..................................................................................................

             3035                                                                            3116
Template     ttgaaagtatttcgatttcttggctttatatatcttgtggaaaggacgaaacaccgactcacgctggatagcctccgtttta
Clonal Se... TTGAAAGTATTTCGATTTCTTGGCTTTATATATCTTGTGGAAAGGACGAAACACCGACTCACGCTGGATAGCCTCCGTTTTA

..................................................................................................

             3117                                                                            3198
Template     gagctagaaatagcaagttaaaataaggctagtccgttatcaacttgaaaaagtggcaccgagtcggtgctttttttcatgg
Clonal Se... GAGCTAGAAATAGCAAGTTAAAATAAGGCTAGTCCGTTATCAACTTGAAAAAGTGGCACCGAGTCGGTGCTTTTTTTCATGG

..................................................................................................

             3199                                                                            3280
Template     tcatagctgtttcctagcggtaaaacgacggccagtgagggcctatttcccatgattccttcatatttgcatatacgataca
Clonal Se... TCATAGCTGTTTCCTAGCGGTAAAACGACGGCCAGTGAGGGCCTATTTCCCATGATTCCTTCATATTTGCATATACGATACA

..................................................................................................

             3281                                                                            3362
Template     aggctgttagagagataattagaattaatttgactgtaaacacaaagatattagtacaaaatacgtgacgtagaaagtaata
Clonal Se... AGGCTGTTAGAGAGATAATTAGAATTAATTTGACTGTAAACACAAAGATATTAGTACAAAATACGTGACGTAGAAAGTAATA

..................................................................................................

             3363                                                                            3444
Template     atttcttgggtagtttgcagttttaaaattatgttttaaaatggactatcatatgcttaccgtaacttgaaagtatttcgat
Clonal Se... ATTTCTTGGGTAGTTTGCAGTTTTAAAATTATGTTTTAAAATGGACTATCATATGCTTACCGTAACTTGAAAGTATTTCGAT

..................................................................................................

             3445                                                                            3526
Template     ttcttggctttatatatcttgtggaaaggacgaaacaccggtcatcttagtcattacctggttttagagctagaaatagcaa
Clonal Se... TTCTTGGCTTTATATATCTTGTGGAAAGGACGAAACACCGGTCATCTTAGTCATTACCTGGTTTTAGAGCTAGAAATAGCAA

..................................................................................................

             3527                                                                            3608
Template     gttaaaataaggctagtccgttatcaacttgaaaaagtggcaccgagtcggtgctttttttcatggtcatagctgtttccta
Clonal Se... GTTAAAATAAGGCTAGTCCGTTATCAACTTGAAAAAGTGGCACCGAGTCGGTGCTTTTTTTCATGGTCATAGCTGTTTCCTA

..................................................................................................
```

```
             3609                                                                          3690
Template     agggtaaaacgacggccagtgagggcctatttcccatgattccttcatatttgcatatacgatacaaggctgttagagagat
Clonal Se... AGGGTAAAACGACGGCCAGTGAGGGCCTATTTCCCATGATTCCTTCATATTTGCATATACGATACAAGGCTGTTAGAGAGAT

...............................................................................................

             3691                                                                          3772
Template     aattagaattaatttgactgtaaacacaaagatattagtacaaaatacgtgacgtagaaagtaataatttcttgggtagttt
Clonal Se... AATTAGAATTAATTTGACTGTAAACACAAAGATATTAGTACAAAATACGTGACGTAGAAAGTAATAATTTCTTGGGTAGTTT

...............................................................................................

             3773                                                                          3854
Template     gcagttttaaaattatgttttaaaatggactatcatatgcttaccgtaacttgaaagtatttcgatttcttggctttatata
Clonal Se... GCAGTTTTAAAATTATGTTTTAAAATGGACTATCATATGCTTACCGTAACTTGAAAGTATTTCGATTTCTTGGCTTTATATA

...............................................................................................

             3855                                                                          3936
Template     tcttgtggaaaggacgaaacaccgggcactgcggctggaggtgggttttagagctagaaatagcaagttaaaataaggctag
Clonal Se... TCTTGTGGAAAGGACGAAACACCGGGCACTGCGGCTGGAGGTGGGTTTTAGAGCTAGAAATAGCAAGTTAAAATAAGGCTAG

...............................................................................................

             3937                                                                          4018
Template     tccgttatcaacttgaaaaagtggcaccgagtcggtgctttttttcatggtcatagctgtttcctcatcgtaaaacgacggc
Clonal Se... TCCGTTATCAACTTGAAAAAGTGGCACCGAGTCGGTGCTTTTTTTCATGGTCATAGCTGTTTCCTCATCGTAAAACGACGGC

...............................................................................................

             4019                                                                          4100
Template     cagtgagggcctatttcccatgattccttcatatttgcatatacgatacaaggctgttagagagataattagaattaatttg
Clonal Se... CAGTGAGGGCCTATTTCCCATGATTCCTTCATATTTGCATATACGATACAAGGCTGTTAGAGAGATAATTAGAATTAATTTG

...............................................................................................

             4101                                                                          4182
Template     actgtaaacacaaagatattagtacaaaatacgtgacgtagaaagtaataatttcttgggtagtttgcagttttaaaattat
Clonal Se... ACTGTAAACACAAAGATATTAGTACAAAATACGTGACGTAGAAAGTAATAATTTCTTGGGTAGTTTGCAGTTTTAAAATTAT

...............................................................................................

             4183                                                                          4264
Template     gttttaaaatggactatcatatgcttaccgtaacttgaaagtatttcgatttcttggctttatatatcttgtggaaaggacg
Clonal Se... GTTTTAAAATGGACTATCATATGCTTACCGTAACTTGAAAGTATTTCGATTTCTTGGCTTTATATATCTTGTGGAAAGGACG

...............................................................................................

             4265                                                                          4346
Template     aaacaccgcacctacctaagaaccatccgttttagagctagaaatagcaagttaaaataaggctagtccgttatcaacttga
Clonal Se... AAACACCGCACCTACCTAAGAACCATCCGTTTTAGAGCTAGAAATAGCAAGTTAAAATAAGGCTAGTCCGTTATCAACTTGA

...............................................................................................

             4347                                                                          4428
Template     aaaagtggcaccgagtcggtgctttttttcatggtcatagctgtttcctacctgtaaaacgacggccagtgagggcctattt
Clonal Se... AAAAGTGGCACCGAGTCGGTGCTTTTTTTCATGGTCATAGCTGTTTCCTACCTGTAAAACGACGGCCAGTGAGGGCCTATTT

...............................................................................................

             4429                                                                          4510
Template     cccatgattccttcatatttgcatatacgatacaaggctgttagagagataattagaattaatttgactgtaaacacaaaga
Clonal Se... CCCATGATTCCTTCATATTTGCATATACGATACAAGGCTGTTAGAGAGATAATTAGAATTAATTTGACTGTAAACACAAAGA

...............................................................................................
```

```
               4511                                                                            4592
Template       tattagtacaaaatacgtgacgtagaaagtaataatttcttgggtagtttgcagttttaaaattatgttttaaaatggacta
Clonal Se...   TATTAGTACAAAATACGTGACGTAGAAAGTAATAATTTCTTGGGTAGTTTGCAGTTTTAAAATTATGTTTTAAAATGGACTA

.................................................................................................

               4593                                                                            4674
Template       tcatatgcttaccgtaacttgaaagtatttcgatttcttggctttatatatcttgtggaaaggacgaaacaccgttcgtatc
Clonal Se...   TCATATGCTTACCGTAACTTGAAAGTATTTCGATTTCTTGGCTTTATATATCTTGTGGAAAGGACGAAACACCGTTCGTATC

.................................................................................................

               4675                                                                            4756
Template       tgtaaaaccaaggttttagagctagaaatagcaagttaaaataaggctagtccgttatcaacttgaaaaagtggcaccgagt
Clonal Se...   TGTAAAACCAAGGTTTTAGAGCTAGAAATAGCAAGTTAAAATAAGGCTAGTCCGTTATCAACTTGAAAAAGTGGCACCGAGT

.................................................................................................

               4757                                                                            4838
Template       cggtgctttttttcatggtcatagctgtttcctgcgagtaaaacgacggccagtgagggcctatttcccatgattccttcat
Clonal Se...   CGGTGCTTTTTTTCATGGTCATAGCTGTTTCCTGCGAGTAAAACGACGGCCAGTGAGGGCCTATTTCCCATGATTCCTTCAT

.................................................................................................

               4839                                                                            4920
Template       atttgcatatacgatacaaggctgttagagagataattagaattaatttgactgtaaacacaaagatattagtacaaaatac
Clonal Se...   ATTTGCATATACGATACAAGGCTGTTAGAGAGATAATTAGAATTAATTTGACTGTAAACACAAAGATATTAGTACAAAATAC

.................................................................................................

               4921                                                                            5002
Template       gtgacgtagaaagtaataatttcttgggtagtttgcagttttaaaattatgttttaaaatggactatcatatgcttaccgta
Clonal Se...   GTGACGTAGAAAGTAATAATTTCTTGGGTAGTTTGCAGTTTTAAAATTATGTTTTAAAATGGACTATCATATGCTTACCGTA

.................................................................................................

               5003                                                                            5084
Template       acttgaaagtatttcgatttcttggctttatatatcttgtggaaaggacgaaacaccgcacggtcaccctgacacgctgttt
Clonal Se...   ACTTGAAAGTATTTCGATTTCTTGGCTTTATATATCTTGTGGAAAGGACGAAACACCGCACGGTCACCCTGACACGCTGTTT

.................................................................................................

               5085                                                                            5166
Template       tagagctagaaatagcaagttaaaataaggctagtccgttatcaacttgaaaaagtggcaccgagtcggtgctttttttcat
Clonal Se...   TAGAGCTAGAAATAGCAAGTTAAAATAAGGCTAGTCCGTTATCAACTTGAAAAAGTGGCACCGAGTCGGTGCTTTTTTTCAT

.................................................................................................

               5167                                                                            5248
Template       ggtcatagctgtttcctcgtagcaagcaagcgctcgaaacggtgcagcggctgttgccggtgctgtgccaggaccatggcct
Clonal Se...   GGTCATAGCTGTTTCCTCGTAGCAAGCAAGCGCTCGAAACGGTGCAGCGGCTGTTGCCGGTGCTGTGCCAGGACCATGGCCT

.................................................................................................

               5249                                                                            5330
Template       gaccccggaccaagtggtggctatcgagacgtctagaccagccaggacagaaatgcctcgacttcgctgctacccaaggttg
Clonal Se...   GACCCCGGACCAAGTGGTGGCTATCGAGACGTCTAGACCAGCCAGGACAGAAATGCCTCGACTTCGCTGCTACCCAAGGTTG

.................................................................................................

               5331                                                                            5412
Template       ccgggtgacgcacaccgtggaaacggatgaaggcacgaacccagtggacataagcctgttcggttcgtaagctgtaatgcaa
Clonal Se...   CCGGGTGACGCACACCGTGGAAACGGATGAAGGCACGAACCCAGTGGACATAAGCCTGTTCGGTTCGTAAGCTGTAATGCAA

.................................................................................................
```

```
             5413                                                                          5494
Template     gtagcgtatgcgctcacgcaactggtccagaaccttgaccgaacgcagcggtggtaacggcgcagtggcggttttcatggct
Clonal Se... GTAGCGTATGCGCTCACGCAACTGGTCCAGAACCTTGACCGAACGCAGCGGTGGTAACGGCGCAGTGGCGGTTTTCATGGCT

...............................................................................................

             5495                                                                          5576
Template     tgttatgactgttttttggggtacagtctatgcctcgggcatccaagcagcaagcgcgttacgccgtgggtcgatgtttga
Clonal Se... TGTTATGACTGTTTTTTGGGGTACAGTCTATGCCTCGGGCATCCAAGCAGCAAGCGCGTTACGCCGTGGGTCGATGTTTGA

...............................................................................................

             5577                                                                          5658
Template     tgttatggagcagcaacgatgttacgcagcagggcagtcgccctaaaacaaagttaaacattatgagggaagcggtgatcgc
Clonal Se... TGTTATGGAGCAGCAACGATGTTACGCAGCAGGGCAGTCGCCCTAAAACAAAGTTAAACATTATGAGGGAAGCGGTGATCGC

...............................................................................................

             5659                                                                          5740
Template     cgaagtatcgactcaactatcagaggtagttggcgtcatcgagcgccatctcgaaccgacgttgctggccgtacatttgtac
Clonal Se... CGAAGTATCGACTCAACTATCAGAGGTAGTTGGCGTCATCGAGCGCCATCTCGAACCGACGTTGCTGGCCGTACATTTGTAC

...............................................................................................

             5741                                                                          5822
Template     ggctccgcagtggatggcggcctgaagccacacagtgatattgatttgctggttacggtgaccgtaaggcttgatgaaacaa
Clonal Se... GGCTCCGCAGTGGATGGCGGCCTGAAGCCACACAGTGATATTGATTTGCTGGTTACGGTGACCGTAAGGCTTGATGAAACAA

...............................................................................................

             5823                                                                          5904
Template     cgcggcgagctttgatcaacgaccttttggaaacttcggcttcccctggagagagcgagattctccgcgctgtagaagtcac
Clonal Se... CGCGGCGAGCTTTGATCAACGACCTTTTGGAAACTTCGGCTTCCCCTGGAGAGAGCGAGATTCTCCGCGCTGTAGAAGTCAC

...............................................................................................

             5905                                                                          5986
Template     cattgttgtgcacgacgacatcattccgtggcgttatccagctaagcgcgaactgcaatttggagaatggcagcgcaatgac
Clonal Se... CATTGTTGTGCACGACGACATCATTCCGTGGCGTTATCCAGCTAAGCGCGAACTGCAATTTGGAGAATGGCAGCGCAATGAC

...............................................................................................

             5987                                                                          6068
Template     attcttgcaggtatcttcgagccagccacgatcgacattgatctggctatcttgctgacaaaagcaagagaacatagcgttg
Clonal Se... ATTCTTGCAGGTATCTTCGAGCCAGCCACGATCGACATTGATCTGGCTATCTTGCTGACAAAAGCAAGAGAACATAGCGTTG

...............................................................................................

             6069                                                                          6150
Template     ccttggtaggtccagcggcggaggaactctttgatccggttcctgaacaggatctatttgaggcgctaaatgaaaccttaac
Clonal Se... CCTTGGTAGGTCCAGCGGCGGAGGAACTCTTTGATCCGGTTCCTGAACAGGATCTATTTGAGGCGCTAAATGAAACCTTAAC

...............................................................................................

             6151                                                                          6232
Template     gctatggaactcgccgcccgactgggctggcgatgagcgaaatgtagtgcttacgttgtcccgcatttggtacagcgcagta
Clonal Se... GCTATGGAACTCGCCGCCCGACTGGGCTGGCGATGAGCGAAATGTAGTGCTTACGTTGTCCCGCATTTGGTACAGCGCAGTA

...............................................................................................

             6233                                                                          6314
Template     accggcaaaatcgcgccgaaggatgtcgctgccgactgggcaatggagcgcctgccggcccagtatcagcccgtcatacttg
Clonal Se... ACCGGCAAAATCGCGCCGAAGGATGTCGCTGCCGACTGGGCAATGGAGCGCCTGCCGGCCCAGTATCAGCCCGTCATACTTG

...............................................................................................
```

```
              6315                                                                              6396
Template      aagctagacaggcttatcttggacaagaagaagatcgcttggcctcgcgcgcagatcagttggaagaatttgtccactacgt
Clonal Se...  AAGCTAGACAGGCTTATCTTGGACAAGAAGAAGATCGCTTGGCCTCGCGCGCAGATCAGTTGGAAGAATTTGTCCACTACGT

...............................................................................................

              6397                                                                              6478
Template      gaaaggcgagatcaccaaggtagtcggcaaataaccctcgagccacccatgaccaaaatcccttaacgtgagttacgcgtcg
Clonal Se...  GAAAGGCGAGATCACCAAGGTAGTCGGCAAATAACCCTCGAGCCACCCATGACCAAAATCCCTTAACGTGAGTTACGCGTCG

...............................................................................................

              6479                                        6522
Template      ttccactgagcgtcagaccccgtagaaaagatcaaaggatcttc
Clonal Se...  TTCCACTGAGCGTCAGACCCCGTAGAAAAGATCAAAGGATCTTC

...............................................................................................
```
