## SupplementaryMaterial for "RAPID-DASH: Fast and Efficient Assembly of Guide RNA Arrays for Multiplexed CRISPR-Cas9 Applications"

**Supplementary Materials**

This document includes the following:

Supplementary Figure 1

Supplementary Figure 2

Supplementary Figure 3

Supplementary Figure 4

Supplementary Table 1

Supplementary Table 2

Supplementary Table 3

Oligos and sequences for gRNA assembly

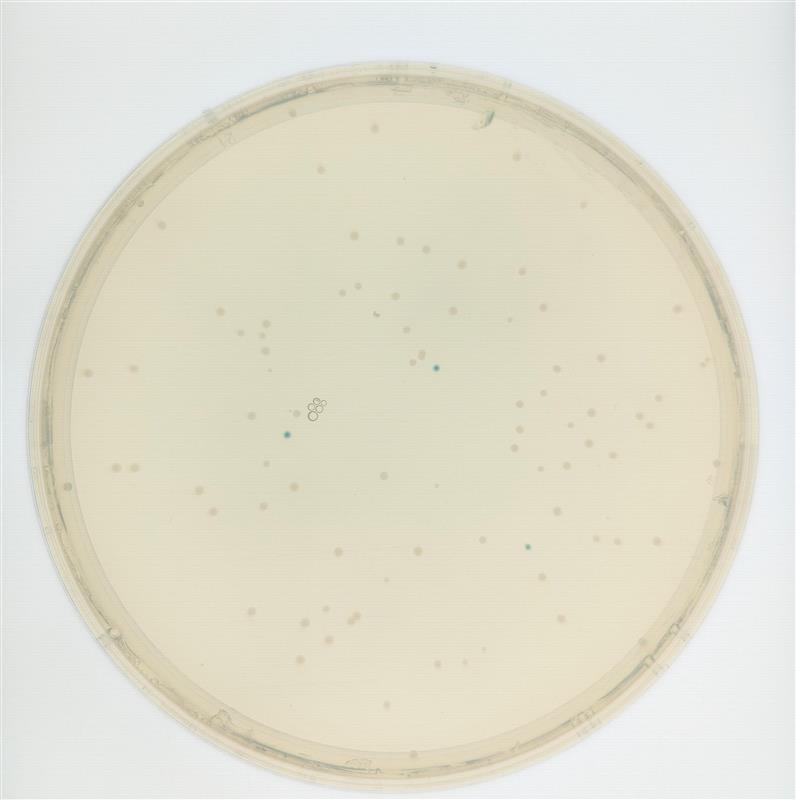

**Supplementary Figure 1:** **Example blue-white screening of bacterial colonies following gRNA array assembly.** White colonies indicate that there was an insertion into the gRNA array cloning site, while blue colonies represent vectors with no insertion.

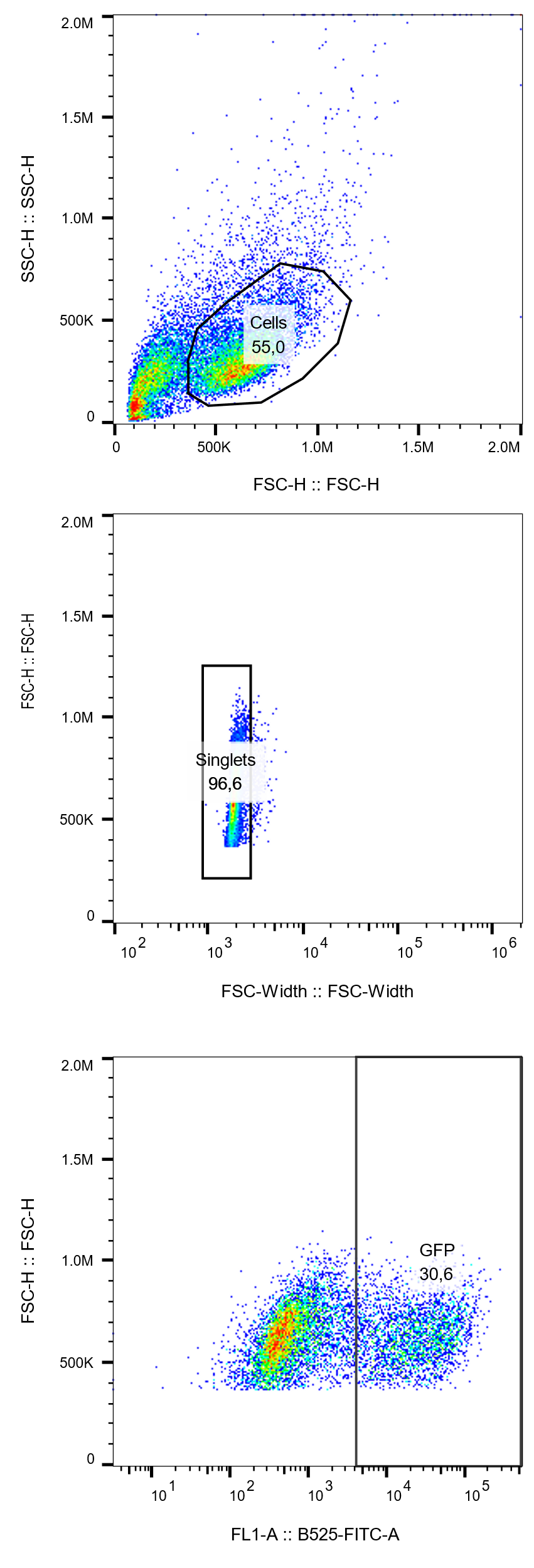

**Supplementary Figure 2: Gating summary for flow cytometry data for GFP reporter activation experiments.** Cells were gated for singlets and then GFP positive cells using FlowJo (v10). Forward vs side scatter height plot was used to gate cells from cellular debris (top), which was then gated for singlets using forward scatter width vs height (middle) and FITC-A channel was used to gate for GFP positive cells from the singlets (bottom).

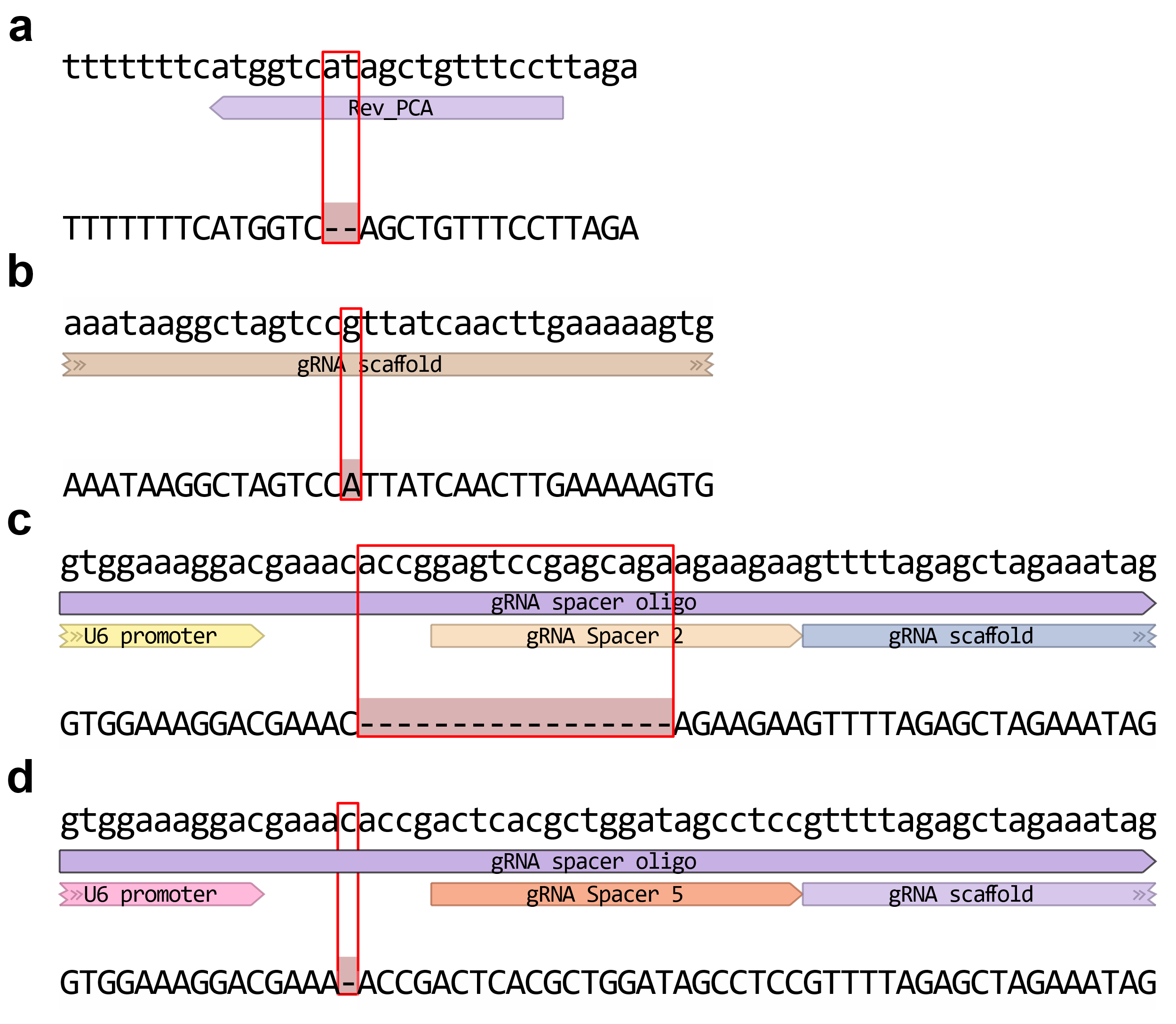

**Supplementary Figure 3: Example mutations seen in gRNA arrays.** Alignments of the consensus sequence of the screened gRNA array clones (bottoms) with desired gRNA assembly plasmid sequence (tops). a) two bp deletion in the priming part of a PCA handle primer. b) 1 bp substitution in the priming part of a PCR handle primer. c) 17 bp deletion and (d) a 1 bp deletion within gRNA spacer oligo

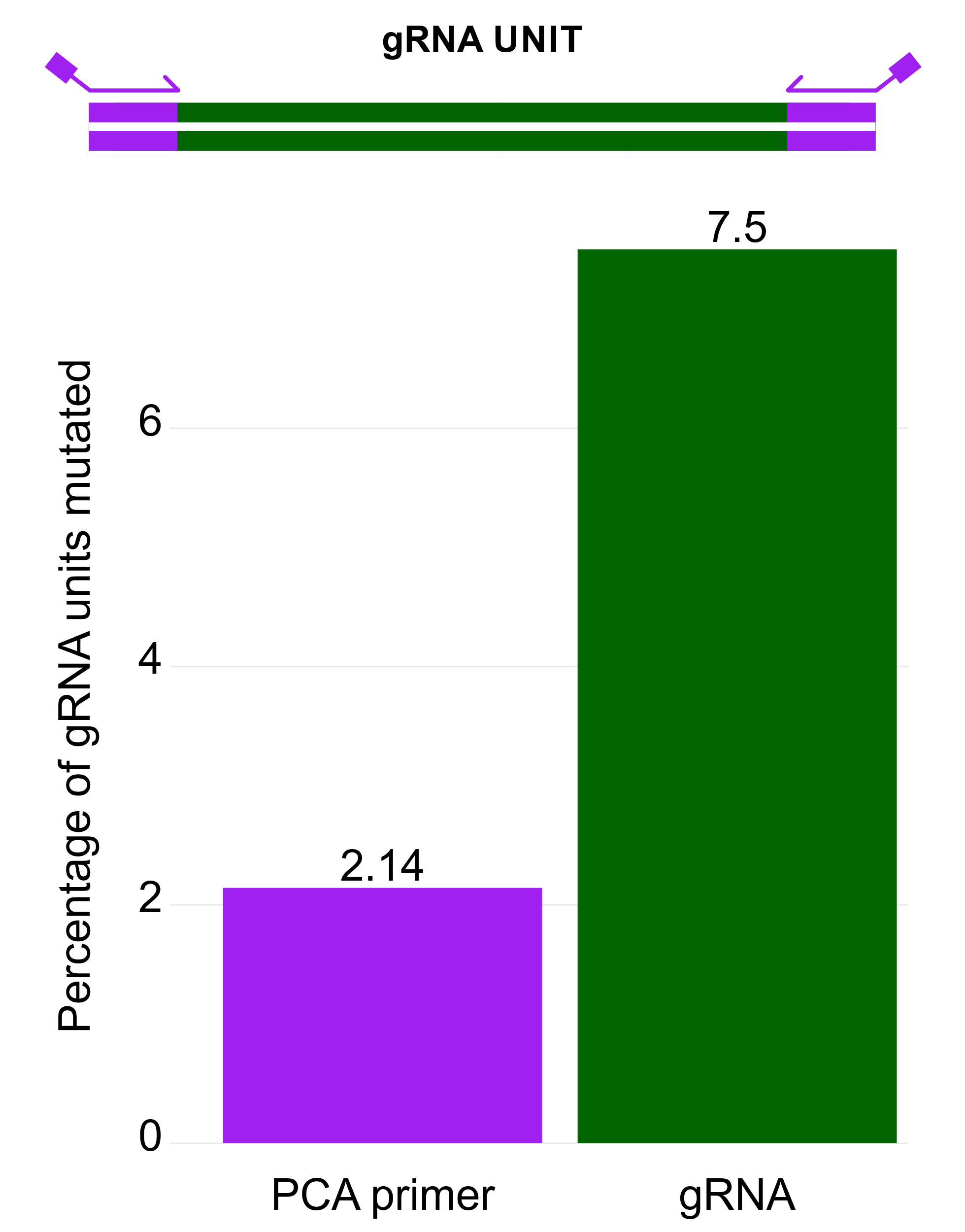

**Supplementary Figure 4:** **Percentage of gRNA units mutated within the screened arrays.** Mutations observed within the gRNA units were categorized based on their occurrence in the PCA primer binding region or the rest of the gRNA units, including the U6 promoter, spacer sequence, and the gRNA scaffold.

**Supplementary Table 1: gRNA array assembly.** Values represent the percentages of extracted gRNA array inserts (from bulk whole plasmid nanopore reads) whose lengths correspond to each number of gRNA units in the array. Full-length 10-gRNA arrays is highlighted in red. “Unclassified” represents inserts that did not match expected array sizes.

| **Number of gRNAs in the array** | **Assembly 1** | **Assembly 2** | **Assembly 3** |
| --- | --- | --- | --- |
| **0** | 0 | 0 | 0 |
| **1** | 3.03 | 0 | 0 |
| **2** | 0 | 0 | 3.68 |
| **3** | 0 | 0 | 2.63 |
| **4** | 1.52 | 0.58 | 4.21 |
| **5** | 0 | 0.58 | 5.79 |
| **6** | 3.03 | 0.58 | 0.53 |
| **7** | 0 | 1.75 | 0 |
| **8** | 4.55 | 1.75 | 1.58 |
| **9** | 1.52 | 1.17 | 3.16 |
| **10** | 81.82 | 87.72 | 74.21 |
| **>10** | 0 | 0.58 | 1.05 |
| **Unclassified** | 4.55 | 5.26 | 3.16 |

**Supplementary Table 2: Read length binning thresholds for calculating the number of gRNAs in an array.** Expected insert lengths were calculated based on the number of gRNAs in an array. Ranges were used to account for high error rate of nanopore sequencing.

| **Number of gRNAs** | **Expected Length (bp)** | **Lower range (bp)** | **Upper Range (bp)** |
| --- | --- | --- | --- |
| >10 | >4311 | 4241 |  |
| 10 | 3917 | 3847 | 3987 |
| 9 | 3523 | 3453 | 3593 |
| 8 | 3129 | 3059 | 3199 |
| 7 | 2735 | 2665 | 2805 |
| 6 | 2341 | 2271 | 2411 |
| 5 | 1947 | 1877 | 2017 |
| 4 | 1553 | 1483 | 1623 |
| 3 | 1159 | 1089 | 1229 |
| 2 | 765 | 695 | 835 |
| 1 | 371 | 301 | 441 |
| 0 | 0 | 0 | 70 |

**Supplementary Table 3:** **Pairwise comparisons of GFP reporter activation for each gRNA position within the array versus non-targeting controls (NTC) and GFP-only controls.** p-values were calculated using two-sample t-tests and adjusted for multiple comparisons using the Bonferroni method within each comparison set (vs. NTC and vs. GFP-only).

| Comparison | p-values | Adjusted p-values | Significance |
| --- | --- | --- | --- |
| G1 vs NTC | 3.66e-06 | 3.66e-05 | *** |
| G2 vs NTC | 4.16e-07 | 4.16e-06 | *** |
| G3 vs NTC | 8.91e-05 | 8.91e-04 | *** |
| G4 vs NTC | 3.67e-04 | 3.67e-03 | ** |
| G5 vs NTC | 6.47e-04 | 6.47e-03 | ** |
| G6 vs NTC | 6.27e-06 | 6.27e-05 | *** |
| G7 vs NTC | 8.11e-04 | 8.11e-03 | ** |
| G8 vs NTC | 1.24e-04 | 1.24e-03 | ** |
| G9 vs NTC | 3.64e-04 | 3.64e-03 | ** |
| G10 vs NTC | 7.72e-04 | 7.72e-03 | ** |
| G1 vs GFP only | 9.74e-03 | 9.74e-02 |  |
| G2 vs GFP only | 4.19e-02 | 4.19e-01 |  |
| G3 vs GFP only | 2.78e-03 | 2.78e-02 | * |
| G4 vs GFP only | 2.29e-03 | 2.29e-02 | * |
| G5 vs GFP only | 7.01e-03 | 7.01e-02 |  |
| G6 vs GFP only | 6.72e-03 | 6.72e-02 |  |
| G7 vs GFP only | 3.57e-03 | 3.57e-02 | * |
| G8 vs GFP only | 2.18e-03 | 2.18e-02 | * |
| G9 vs GFP only | 2.45e-03 | 2.45e-02 | * |
| G10 vs GFP only | 3.26e-03 | 3.26e-02 | * |

Statistical significance is marked with * as follows: * - adjusted p-value < 0.05, ** - adjusted p-value < 0.01 and *** - adjusted p-value < 0.001.

### Oligos and sequences for gRNA assembly

**Note:** Based on our finding that the primers are a primary cause of mutations in final gRNA array assemblies, the forward and reverse primers for the initial amplification of the U6 promoter and gRNA terminator scaffold are one of the only places that we think makes economic sense to purify (e.g. HPLC to increase the purity of full-length oligos). This is because the PCR products they produce are used in every single gRNA unit and they could introduce mutations that adversely affect gRNA function. The gRNA unit amplification oligos are priming in inconsequential regions and so any introduced mutations are likely benign, and gRNA spacer oligos are often used only once and so it may or may not be economical to purify them. The results in this study reflect no purification of any oligos.

#### U6 Promoter

**Forward primer**: GTAAAACGACGGCCAGTgagggcctatttcccatgattc

**Reverse primer**: GGTGTTTCGTCCTTTCCAC

**Amplicon sequence**: GTAAAACGACGGCCAGTgagggcctatttcccatgattccttcatatttgcatatacgatacaaggctgttagagagataattggaattaatttgactgtaaacacaaagatattagtacaaaatacgtgacgtagaaagtaataatttcttgggtagtttgcagttttaaaattatgttttaaaatggactatcatatgcttaccgtaacttgaaagtatttcgatttcttggctttatatatcttGTGGAAAGGACGAAACACC

#### gRNA terminator scaffold

**Forward primer**: gttttagagctaGAAAtagcaag

**Reverse primer**: AGGAAACAGCTATGACCATGAAAAAAAgcaccgactcggtgccac

**Amplicon sequence**: gttttagagctaGAAAtagcaagttaaaataaggctagtccgttatcaacttgaaaaagtggcaccgagtcggtgcTTTTTTTCATGGTCATAGCTGTTTCCT

#### gRNA spacer oligos

**Sample**: GTGGAAAGGACGAAACACCgNNNNNNNNNNNNNNNNNNNNgttttagagctaGAAAtag

|  | **gRNA spacer** | **Ordered oligo** |
| --- | --- | --- |
| 1 | GGAATCCCTTCTGCAGCACC | GTGGAAAGGACGAAACACCgGGAATCCCTTCTGCAGCACCgttttagagctaGAAAtag |
| 2 | GAGTCCGAGCAGAAGAAGAA | GTGGAAAGGACGAAACACCgGAGTCCGAGCAGAAGAAGAAgttttagagctaGAAAtag |
| 3 | TTTATCACAGGCTCCAGGAA | GTGGAAAGGACGAAACACCgTTTATCACAGGCTCCAGGAAgttttagagctaGAAAtag |
| 4 | GGCCCAGACTGAGCACGTGA | GTGGAAAGGACGAAACACCgGGCCCAGACTGAGCACGTGAgttttagagctaGAAAtag |
| 5 | ACTCACGCTGGATAGCCTCC | GTGGAAAGGACGAAACACCgACTCACGCTGGATAGCCTCCgttttagagctaGAAAtag |
| 6 | GTCATCTTAGTCATTACCTG | GTGGAAAGGACGAAACACCgGTCATCTTAGTCATTACCTGgttttagagctaGAAAtag |
| 7 | GGCACTGCGGCTGGAGGTGG | GTGGAAAGGACGAAACACCgGGCACTGCGGCTGGAGGTGGgttttagagctaGAAAtag |
| 8 | CACCTACCTAAGAACCATCC | GTGGAAAGGACGAAACACCgCACCTACCTAAGAACCATCCgttttagagctaGAAAtag |
| 9 | TTCGTATCTGTAAAACCAAG | GTGGAAAGGACGAAACACCgTTCGTATCTGTAAAACCAAGgttttagagctaGAAAtag |
| 10 | GTATCTAGTGTTGGTGTCCT | GTGGAAAGGACGAAACACCgGTATCTAGTGTTGGTGTCCTgttttagagctaGAAAtag |
| GFP targeting | CACGGTCACCCTGACACGCT | GTGGAAAGGACGAAACACCgCACGGTCACCCTGACACGCTgttttagagctaGAAAtag |

#### Primers for gRNA unit amplification

| **gRNA unit** | **Primer direction** | **Sequence** | **BsaI Overhang** |
| --- | --- | --- | --- |
| **1** | **Fwd** | ATAAGGATCCGGTCTCAGGTAGTAAAACGACGGCCAGT | GGTA |
| **1** | **Rev** | ATAATGTACAGGTCTCTTCTAAGGAAACAGCTATGACCATG | TCTA |
| **2** | **Fwd** | ATAAGGATCCGGTCTCATAGAGTAAAACGACGGCCAGT | TAGA |
| **2** | **Rev** | ATAATGTACAGGTCTCTGGAGAGGAAACAGCTATGACCATG | GGAG |
| **3** | **Fwd** | ATAAGGATCCGGTCTCACTCCGTAAAACGACGGCCAGT | CTCC |
| **3** | **Rev** | ATAATGTACAGGTCTCTTGATAGGAAACAGCTATGACCATG | TGAT |
| **4** | **Fwd** | ATAAGGATCCGGTCTCAATCAGTAAAACGACGGCCAGT | ATCA |
| **4** | **Rev** | ATAATGTACAGGTCTCTTCAGAGGAAACAGCTATGACCATG | TCAG |
| **5** | **Fwd** | ATAAGGATCCGGTCTCACTGAGTAAAACGACGGCCAGT | CTGA |
| **5** | **Rev** | ATAATGTACAGGTCTCTCGCTAGGAAACAGCTATGACCATG | CGCT |
| **6** | **Fwd** | ATAAGGATCCGGTCTCAAGCGGTAAAACGACGGCCAGT | AGCG |
| **6** | **Rev** | ATAATGTACAGGTCTCTCCTTAGGAAACAGCTATGACCATG | CCTT |
| **7** | **Fwd** | ATAAGGATCCGGTCTCAAAGGGTAAAACGACGGCCAGT | AAGG |
| **7** | **Rev** | ATAATGTACAGGTCTCTGATGAGGAAACAGCTATGACCATG | GATG |
| **8** | **Fwd** | ATAAGGATCCGGTCTCACATCGTAAAACGACGGCCAGT | CATC |
| **8** | **Rev** | ATAATGTACAGGTCTCTAGGTAGGAAACAGCTATGACCATG | AGGT |
| **9** | **Fwd** | ATAAGGATCCGGTCTCAACCTGTAAAACGACGGCCAGT | ACCT |
| **9** | **Rev** | ATAATGTACAGGTCTCTTCGCAGGAAACAGCTATGACCATG | TCGC |
| **10** | **Fwd** | ATAAGGATCCGGTCTCAGCGAGTAAAACGACGGCCAGT | GCGA |
| **10** | **Rev** | ATAATGTACAGGTCTCTTACGAGGAAACAGCTATGACCATG | TACG |
